# A transient brain endothelial translatome response to endotoxin is associated with mild cognitive changes post-shock in young mice

**DOI:** 10.1101/2024.03.03.583191

**Authors:** Shuhan Lu, Iria Di John Portela, Nina Martino, Ramon Bossardi Ramos, Abigail E Salinero, Rachel M Smith, Kristen L Zuloaga, Alejandro P Adam

## Abstract

Sepsis-associated encephalopathy (SAE) is a common manifestation in septic patients that is associated with increased risk of long-term cognitive impairment. SAE is driven, at least in part, by brain endothelial dysfunction in response to systemic cytokine signaling. However, the mechanisms driving SAE and its consequences remain largely unknown. Here, we performed translating ribosome affinity purification and RNA-sequencing (TRAP-seq) from the brain endothelium to determine the transcriptional changes after an acute endotoxemic (LPS) challenge. LPS induced a strong acute transcriptional response in the brain endothelium that partially correlates with the whole brain transcriptional response and suggested an endothelial-specific hypoxia response. Consistent with a crucial role for IL-6, loss of the main regulator of this pathway, SOCS3, leads to a broadening of the population of genes responsive to LPS, suggesting that an overactivation of the IL-6/JAK/STAT3 pathway leads to an increased transcriptional response that could explain our prior findings of severe brain injury in these mice. To identify any potential sequelae of this acute response, we performed brain TRAP-seq following a battery of behavioral tests in mice after apparent recovery. We found that the transcriptional response returns to baseline within days post-challenge. Despite the transient nature of the response, we observed that mice that recovered from the endotoxemic shock showed mild, sex-dependent cognitive impairment, suggesting that the acute brain injury led to sustained, non-transcriptional effects. A better understanding of the transcriptional and non-transcriptional changes in response to shock is needed in order to prevent and/or revert the devastating consequences of septic shock.

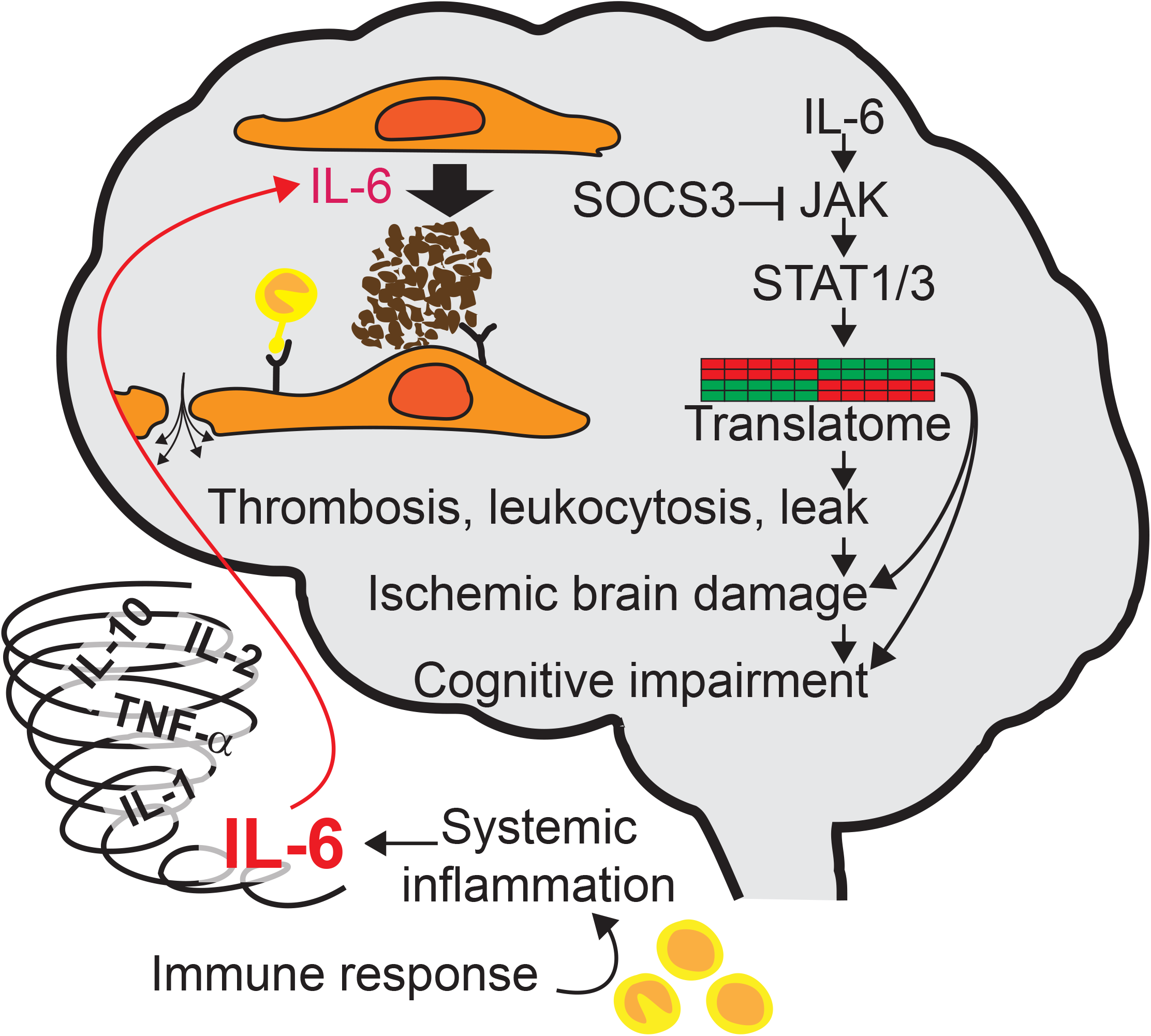

## Introduction

Sepsis-associated encephalopathy (SAE) is a common manifestation in septic patients. It is characterized by diffuse cerebral dysfunction in the absence of a direct brain infection and often clinically manifests as delirium or coma (Fong TG et al., 2009;Gofton TE and Young GB, 2012;Hughes CG et al., 2016;Tokuda R et al., 2023). Acute encephalopathy occurs approximately in 70% of patients admitted at intensive care units (ICU) (Mazeraud A et al., 2020). Delirium usually resolves with the patient’s overall improvement (Fong TG, et al., 2009). However, occurrence of SAE is associated with increased ICU stay, six-month mortality post-discharge, and higher risk of long-term cognitive impairment and dementia (Gofton TE and Young GB, 2012;Iwashyna TJ et al., 2010). The risks are higher in patients over 65 years old (Peterson JF et al., 2006) and those with more severe sepsis (Tokuda R, et al., 2023). Patients admitted at the ICU due to shock are at high risk of developing multiorgan failure, in large part due to a dysregulated inflammatory reaction that involves large concentrations of circulating cytokines, a condition usually known as a ‘cytokine storm’ (Tisoncik JR et al., 2012). The mechanisms driving encephalopathy are thought to involve an impaired microvascular blood-brain barrier, coagulopathy, and reduced perfusion, in response to systemic inflammatory cytokines (Crippa IA et al., 2023;Daneman R and Prat A, 2015;Gofton TE and Young GB, 2012;Hughes CG, et al., 2016;Peng X et al., 2021;Schramm P et al., 2012;Sweeney MD et al., 2019). Thus, targeting the brain microvasculature response to shock is a promising avenue to prevent or limit SAE. However, we do not fully understand the mechanisms by which the brain endothelium responds to severe systemic inflammatory signals that lead to brain microvascular dysfunction and SAE.

Thse role of interleukin-6 (IL-6) in systemic inflammatory conditions is well established (Hou T et al., 2015;Hunter CA and Jones SA, 2015;Remick DG et al., 2005). Prior work showed that IL-6 promotes a sustained loss of endothelial barrier function, both directly (Alsaffar H et al., 2018) and as part of an autocrine loop in response to LPS or TNF (Martino N et al., 2022). We previously reported that a single LPS injection led to increased brain vascular permeability 15 hours post-challenge (Martino N et al., 2021). Mice lacking endothelial expression of SOCS3 (SOCS3iEKO), the main negative regulator of the IL-6/JAK/STAT3 pathway, displayed significantly higher brain microvascular leak and organ injury than wild-type (WT) control mice (Martino N, et al., 2021), demonstrating a critical role for this pathway in the regulation of the blood-brain barrier in response to systemic inflammation.

The mechanisms driving LPS- and IL-6-induced brain endothelial dysfunction and its consequences, however, remain largely unknown. To begin addressing this mechanism, we sought to perform translating ribosome affinity purification (TRAP) (Zhou P et al., 2013) and RNA-sequencing (TRAP-seq) from the brain endothelium to determine the transcriptional changes after an acute challenge. To determine potential sequelae, we performed TRAP-seq following a battery of behavioral tests in mice after apparent recovery. We found that LPS induces a strong acute transcriptional response in the brain endothelium that returns to baseline within days post-challenge. Consistent with a critical role for the IL-6 pathway, SOCS3iEKO mice displayed an exacerbated transcriptional response. Despite the transient transcriptional changes, mice displayed mild, sex-specific cognitive impairment, suggesting lasting effects in brains exposed to a single endotoxemic shock.

## Experimental Procedures

### Materials, antibodies, and primers

The commercial sources for critical reagents and their catalog numbers are listed in Supplemental Table 1. Supplemental Table 2 lists all primers used.

### Mice

All mouse experiments were approved by the Animal Resource Facility (ARF) at Albany Medical College and were performed in accordance with the Institutional Animal Care and Use Committee (IACUC)-approved protocols and followed MQTiPSS(Osuchowski MF et al., 2018) guidelines for randomization, masking, and euthanasia following standardized criteria.

Endothelial-specific TRAP mice generation was recently reported (Bossardi Ramos R et al., 2023). Briefly, B6.Tg(Cdh5-cre/ERT2)1Rha (Cdh5-CreERT2 endothelial driver) mice (Wang Y et al., 2010) were crossed with B6.129S4-Gt(ROSA)26Sortm1(CAG-EGFP/Rpl10a,-birA)Wtp/J (Rosa26fsTRAP) mice (Zhou P, et al., 2013) previously backcrossed to a C57Bl/6 background (a kind gift from Dr Patrick Murphy, University of Connecticut) and with B6;129S4-Socs3tm1Ayos/J (SOCS3^fl/fl^ conditional knockout) (Yasukawa H et al., 2003) mice (The Jackson Laboratory). Tamoxifen-inducible endothelial-specific SOCS3 KO and WT littermate control mice for all experiments were generated by crossing two SOCS3^fl/+^ heterozygote mice. All experimental mice carried homozygous Cre and GFP/Rpl10a transgenes.

Genotypes and sex were confirmed with PCR genotyping (Supplemental Table 2). All mice received IP injections of 2mg/100μl tamoxifen for 5 consecutive days. All the experiments were performed 2 to 3 weeks after the end of tamoxifen injections. Mice were housed in pathogen-free rooms with controlled humidity and temperature, and 12 h light/dark cycles. A group of equal to or less than five mice lived in Allentown cages with accessible food and water ad libitum.

Acute systemic inflammation was induced by a single IP injection of 250μg LPS dissolved in 250μl of sterile saline solution. Control mice were injected with 250μl of sterile saline solution. Severity scores were assessed 13 hours after the LPS injection based on a scoring system as previously described (Bossardi Ramos R, et al., 2023;Martino N, et al., 2021). Immediately after, weight and temperature were measured. Mice were either euthanized 15 hours after the LPS injection, or scored (severity, temperature, weight) daily for four days. Behavioral tests as described below were performed between days five and eight. Following euthanasia, organs were collected for TRAP-seq.

### TRAP-seq and bioinformatics

Endothelial RNA isolation was performed with modified established TRAP assay protocols (Heiman M et al., 2014;Zhou P, et al., 2013). Mice were euthanized with an overdose of pentobarbital. The chest cavity was opened immediately after confirmation of the absence of a paw reflex, and mice were subjected to an intracardiac perfusion with ice-cold HBSS containing 100 μg/ml CHX for 2 minutes at a rate of 5 ml/min. Brains were immediately removed and placed in plates containing 1 ml of dissection buffer and minced with razor blades pre-rinsed with RNAse Away solution. Approximately 200 mg of minced tissue was moved to RNAse-free tubes and lysed using RNAse-free disposable pestles.

Samples were then centrifuged at 4 °C, 2000x g for 10 minutes. The supernatant was removed to a new tube and then 1/9 volume of 10% Igepal and 1/9 volume of 300 mM DHPC were added. Tubes were mixed by inversion several times and incubated for 5 minutes on ice. Then, samples were centrifuged at 4 °C 20,000x g for 10 minutes. 50 μl of the supernatant was transferred to a new tube for processing total RNA and the rest was transferred to another new tube for ribosome pulldown for endothelial-specific RNA. For total RNA, 500 μl of Trizol were added to each tube and RNA extraction was conducted with 100 μl of Chloroform. Then supernatant was moved to RNeasy Micro spin columns and an equal volume of 70% EtOH (approx. 350μl) was added to purify RNA by following the manufacturer’s instructions. For the TRAP RNA, supernatants were incubated with prewashed Dynabeads on a nutating rocker for 15 minutes at RT to remove non-specific binding to the beads, and magnets were used for separation. The supernatant was then incubated with Dynabeads bound to anti-EGFP antibodies on a nutating rocker for 45 minutes at 4 °C. Followed by the washes with RNA buffer, the beads reached RT and then incubated for an extra 5 minutes in RLT buffer containing β-mercaptoethanol. RNA was then purified using RNeasy Micro spin columns following the manufacturer’s instructions.

A total of 500 ng (whole brain) or 100 ng (brain TRAP) RNA from each brain was sent to Azenta Life Sciences for standard (bulk RNA-seq) or ultra-low RNA sequencing (TRAP-seq). Raw sequencing files were deposited in NCBI’s Gene Expression Omnibus and are accessible through GEO Series accession number GSE253438. Mapping to the mouse genome (mm10) was performed in R/Bioconductor (Huber W et al., 2015) using the Rsubread (Liao Y et al., 2019) package. Differential gene expression was calculated using edgeR (Robinson MD et al., 2010) package. Gene Set Enrichment Analysis of Gene Ontology was performed using ClusterProfiler (Yu G et al., 2012) package. Upstream analysis (enrichment in pathways and transcription factors) of each datasets was performed using the DecoupleR (Badia IMP et al., 2022) package, the PROGENy model (Schubert M et al., 2018), and the CollecTRI network (Muller-Dott S et al., 2023).

### Behavioral tests

Except for the nest building assay, mice were moved to a room with dim light for 1 h to acclimate to the surrounding. The maze and arena were cleaned with 70% ethanol between trials. After each test, mice were moved to a “recovery cage” to separate the mouse from all untested cage mates. Mouse performances were recorded with a camera placed above the behavioral maze or arena, and mouse movement was tracked and analyzed with ANY-maze software.

Y-maze: Mice were placed in a Y-shaped maze with 3 arms for 5min to freely explore the routes.

Data was analyzed manually by counting the number of alternating entries to each of the three arms. The percent alternation index was calculated as [triad/(number of arm entries-2)]*100. Mice that jumped out of the maze were excluded.

Open field: Mice were placed individually in the arena (box 495 x495 mm) for 10 min. Mouse activities were recorded with the video camera on top of the box and mouse movements were tracked with ANY-maze software. A 4×4 grid was drawn over the video to track the animal’s position over time. The percent center time index was obtained by dividing the time spent in the 4 center squares over the total experimental time.

Novel object recognition (NOR) test: Mice were placed individually in the same arena as for the open field test but containing two identical rubber ducks 8 cm from the wall in the northeast corner and northwest corner of the arena respectively. Training was performed by placing the mice for 10 minutes in the arena to familiarize the mice with these objects. Mice were then returned to their cages. After 1h, mice were returned to the arena for the test trial. At this time, the right rubber duck was replaced with a glass cup of a similar size. Mouse movements were tracked with ANY-maze software for 10 minutes. The time spent on each object was evaluated by determining the time spent within a 5 cm radius from the object. The recognition index was calculated as [(time with novel object / total time with objects)*100].

Nest building: Following the NOR test, mice were single-caged with two nestlets and allowed to freely make nests overnight (16 h). Mice were then removed back to their original group cages for later tissue collection. Nests were scored from 1-5 with 0.5 increments, according to a well-established scale (Deacon RM, 2006). Briefly, scores were: 1, >90% intact nestlet; 2, partially torn nestlet; 3, nestlet mostly shredded but often no identifiable nest site; 4, an identifiable but flat nest; 5, a near perfect nest. Scores from three independent scorers were averaged.

### Cell culture

HCMEC/D3 were obtained from Millipore Sigma (catalog number SCC066). HBEC-5i cells were purchased from ATCC (catalog #CRL-3245) and maintained following manufacturer’s instructions.

HCMEC/D3 cells were cultured in EndoGRO-MV Complete Culture Media (Millipore Sigma). HBEC-5i cells were cultured in DMEM:F-12 1:1 mixture with L-glutamine and HEPES media (ATCC 30-2006TM) containing 40g/mL endothelial growth supplement (ECGS) and 10% fetal bovine serum. To induce IL-6 signaling, cells were treated with a combination of 200 ng/mL recombinant human IL-6, 2 μg/mL LPS, or 10 ng/mL TNF in the presence or absence of 100 ng/mL sIL-6Rα (sIL6R). A similar amount of PBS was used for controls. HUVEC were isolated and grown as described previously (Bossardi Ramos R, et al., 2023;Martino N, et al., 2022;Martino N, et al., 2021).

### RT-qPCR

After cell treatments, cells were lysed with Trizol and total RNA was isolated according to manufacturer’s instructions. Primescript RT Master Mix was used to make cDNA from 400ng of total RNA following manufacturer’s instructions. cDNA was diluted 10-fold in nuclease-free water. 2 µL of cDNA was used per PCR reaction. qPCR was performed in a StepOnePlus or a QuantStudio 3 (Applied Biosystems) instrument using SYBR green-based iTaq supermix (Bio-Rad) and 2 pmol primers (Thermo Fisher). Fold induction was calculated using the ΔΔCt method using GAPDH as housekeeping gene. Primer sequences utilized are listed in Supplemental Table 2.

### Statistics

All statistical analysis and graphs were made in GraphPad Prism, version 6. Analysis for RNA expression levels was performed using a one-way ANOVA comparing all samples versus a control (designated in each figure). Analysis for behavioral tests was performed using a two-way ANOVA and Holm–Šídák post hoc tests comparing all samples versus a control group (designated in each figure). A two-tailed *P* value of less than 0.05 was considered significant.

## Results

### Systemic inflammation induces an acute proinflammatory brain microvascular transcriptional response

To study brain endothelial-specific transcriptional response to systemic inflammation, we crossed Rosa26 fsTRAP transgene with mice carrying a cdh5-CreERT2 inducible endothelial Cre driver. After tamoxifen treatment, mice were challenged with a single intraperitoneal bolus of LPS or vehicle (saline solution). Fifteen hours later, mice were euthanized and whole brains were processed for bulk RNA-seq (whole brain gene expression) and TRAP-seq (endothelial-specific translatome) (Figure 1A). A strong enrichment in endothelial-specific markers and a loss of non-endothelial gene expression demonstrate the efficiency and specificity of the TRAP protocol (Figure 1B). The TRAP-seq response is consistent with endothelial responses measured by single-cell RNA-seq in recent studies of aging and neuroinflammation (Allen WE et al., 2023;Jeong HW et al., 2022). We then compared our data with published brain TRAP from Tie2-Cre;Rpl22-HA mice (Cleuren ACA et al., 2019). Consistent with the increased specificity of the cdh5-CreERT2 (Payne S et al., 2018), we found that many genes induced by LPS in the Tie2-Cre-driven TRAP were not altered in the cdh5-CreERT2-driven TRAP data (Figure 1C). This is likely not due to differences in the LPS challenge, because the total brain RNA-seq demonstrated a strong correlation between the two datasets (Figure 1D).

**Figure 1.**
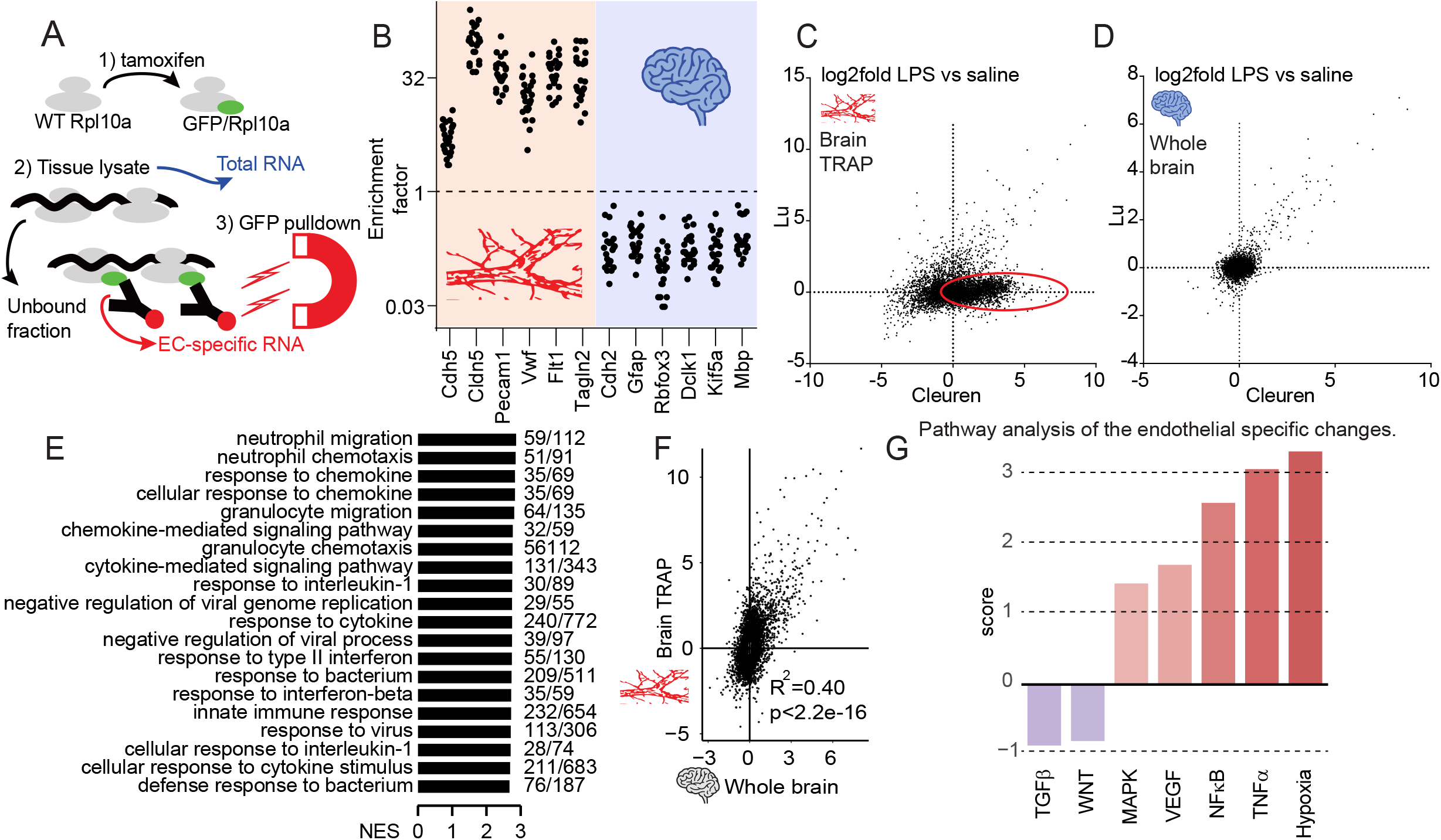
Transcriptional response to LPS in the brain endothelium. (A) Schematic of TRAP approach. (B) Ratio of TRAP-seq over whole brain RNA-seq normalized counts obtained from the same brains. (C) Scatter plots to illustrate the extent of correlation between the brain endothelial responses to LPS as measured in this manuscript (Lu) or by a Tie2-Cre driver (Cleuren ACA, et al., 2019). The red ellipse denotes the gene set responsive in tie2-Cre+ cells but not in cdh5-CreERT2+ mice. (D) Same as in C, but data reflects bulk brain RNA-Seq demonstrating strong identity between the two datasets. (E) Top 20 entries (by normalized enrichment score, NES) of the gene set enrichment analysis of the endothelial transcriptional response to LPS. Shown at the right of each category are the number of enriched genes in the dataset and the total number of genes for each category. (F) Plot showing the correlation between the LPS response (log2 fold change) in whole brain vs brain TRAP. (G) Upstream pathway analysis of the endothelial-specific response, defined as the gene set that is significantly altered by LPS (fold change <0.5 or > 2, adj. p value < 0.05) in the brain TRAP-seq dataset but not in the whole brain bulk RNA-seq dataset.

As expected, gene set enrichment analysis of the endothelial transcriptional response to LPS demonstrates a strong activation of innate immunity and proinflammatory signaling (Figure 1E). This challenge induced strong neuroinflammation, as determined by increased expression of GFAP, TNF, and many other proinflammatory markers in whole brain (Supplemental Table 3). Although the transcriptional endothelial response to LPS showed a strong correlation with that of the whole brain (Figure 1F), multiple genes altered only in the TRAP-seq dataset suggested a marked endothelial-specific hypoxic response (Figure 1G).

We then compared the transcriptional response of SOCS3iEKO mice to littermate control mice.

SOCS3 deletion on SOCS3iEKO mice was confirmed by the TRAP-seq data (Figure 2A). The highest responsive genes to LPS were only modestly affected by the loss of SOCS3 (Figure 2B), suggesting an already maximal response to this gene subset. However, SOCS3iEKO mice responded with a much wider transcriptional response to LPS (Figure 2C-D, Supplemental Table 4). Pathway and transcription factor enrichment analysis of the SOCS3iEKO response demonstrated a strong activation of the JAK/STAT pathway and other transcription factors we previously identified as regulated by IL-6 (Figure 2E-G).

**Figure 2.**
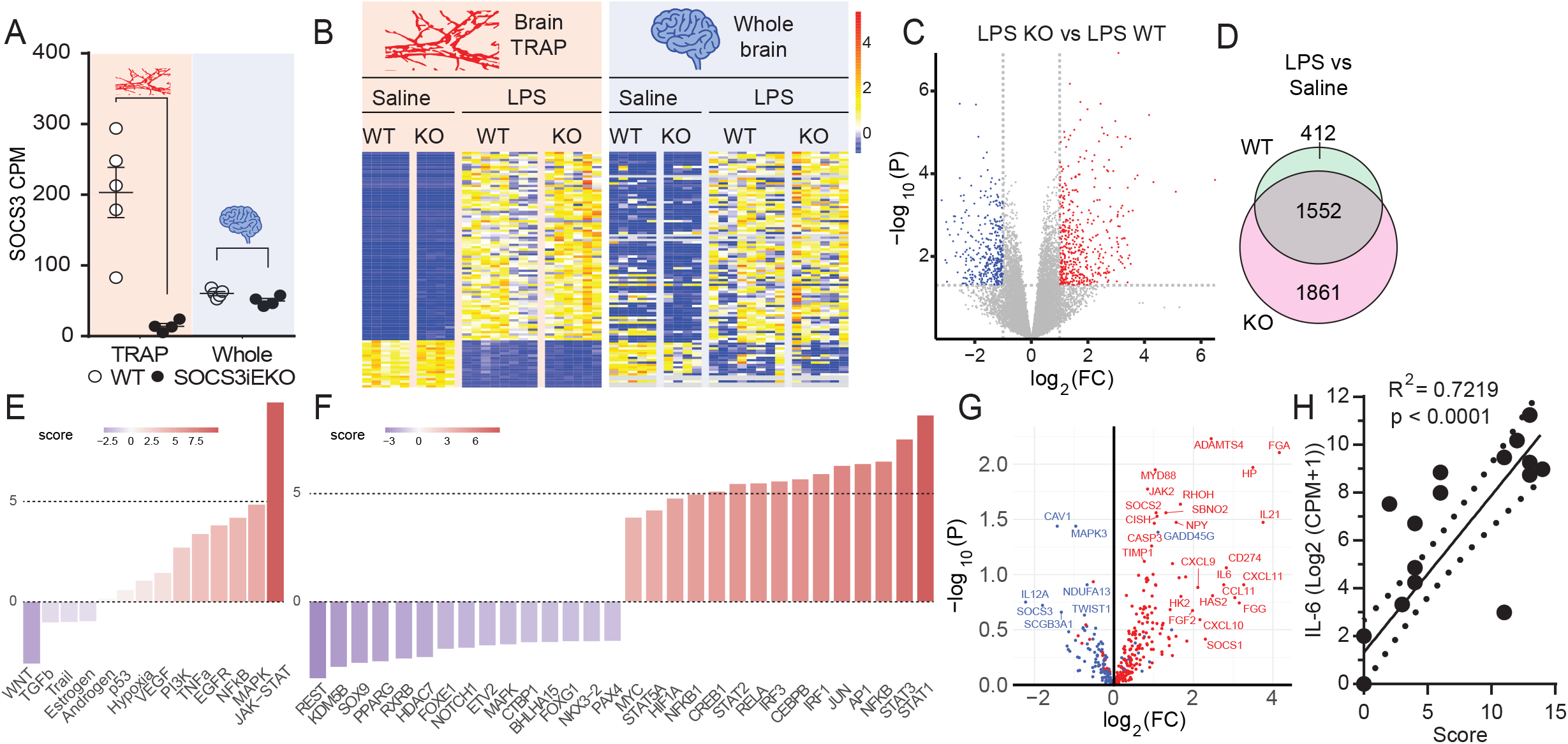
Transcriptional dependence on the IL-6/STAT3/SOCS3 signaling axis. (A) Normalized counts for SOCS3 in brain TRAP and whole brain datasets from saline-treated mice. (B) Heatmap of the 100 most significantly changed genes in the brain endothelium (left) and the corresponding changes in the whole brain (right) of WT and SOCS3iEKO mice. Genes in gray were not detected by whole brain RNA-seq. (C) Volcano plot of the differential expression in response to LPS for either WT or SOCS3iEKO mice. (D) Venn diagram describing the number of differentially expressed genes in response to LPS (right). (E) Upstream pathway analysis of this dataset demonstrating the overactivation of JAK/STAT signaling. (F) Transcription factor enrichment confirming STAT1/3 activity. (G) Volcano plot of log10 p value vs log2 fold change (LPS-treated SOCS3iEKO vs WT mice) for all STAT3-dependent genes (as determined by DecoupleR/CollecTRI). (H) Correlation of endothelial levels of IL-6 gene expression vs severity score.

Consistent with a crucial role for IL-6 driving organ damage, the level of expression in the brain endothelium correlated with the disease severity scoring (Figure 2H).

### The proinflammatory response quickly returns to baseline levels upon shock resolution

We then sought to determine if this acute LPS challenge led to behavioral deficits. We tracked the disease severity (Bossardi Ramos R, et al., 2023;Martino N, et al., 2021) for four days prior to a battery of behavioral tests and brain TRAP-seq (Figure 3A). We performed this assay only in WT mice, since SOCS3iEKO mice do not survive this LPS challenge (Martino N, et al., 2021). All mice had completely recovered by day 4 (Figure 3B). Brain TRAP-Seq and whole brain RNA-seq data from mice after recovery from LPS demonstrate the transient nature of the majority of the endothelial transcriptional responses (Figure 3C, Supplemental Table 5). However, a limited inflammatory response to LPS remains, with increased brain endothelial mRNA levels of several proinflammatory genes (Supplemental Table 5). Consistent with the transient nature of the inflammatory response, we detected limited, sex-dependent behavioral changes in recovering mice compared to saline-treated mice. We found that female mice treated with LPS performed poorly in the nest building test (Figure 3D), while LPS-treated males displayed deficits in the Y maze test (Figure 3E). Both males and females showed no changes in the novel object recognition and the open field tests (Figure 3F-G). No sex-dependent baseline differences were observed in saline-treated mice (Figure 3D-G).

**Figure 3.**
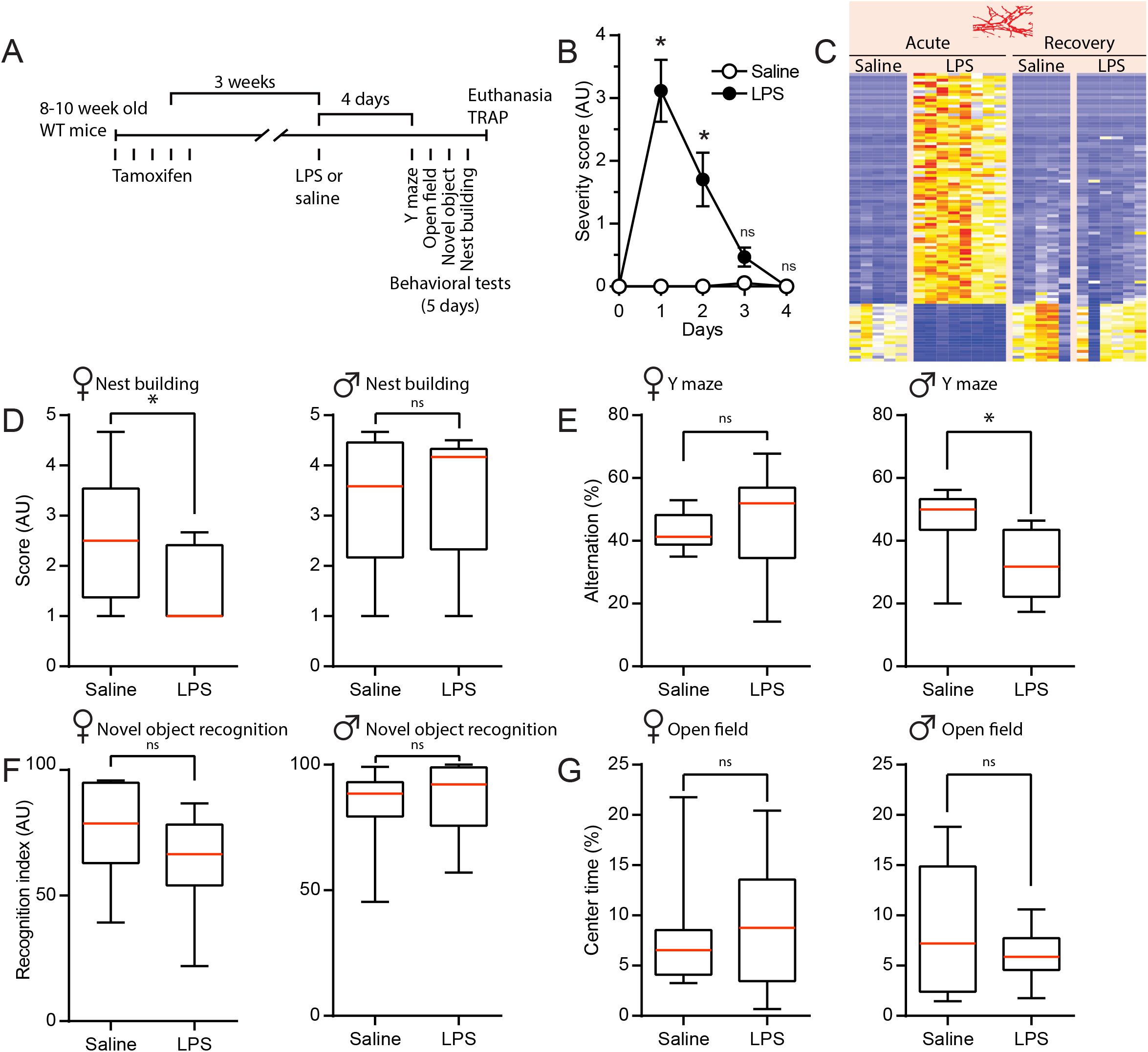
Sustained cognitive impairment despite transitory transcriptional changes. (A) Diagram of the experimental timeline. (B) Daily severity score after saline or LPS injection. Two-way ANOVA of repeated measurements. (C) Heatmap of the top 100 most significantly changed genes 15 hours post saline or LPS injection (left) and the corresponding genes 8 days after saline or LPS injection (right). (D-G) Behavioral tests following recovery from LPS. Assays for nest building (D), Y-maze (E), novel object recognition (F) and open field (G) were performed according to the timeline shown in A. Mann Whitney test (n=7—10 per group). Asterisks, p < 0.05.

### Endothelial cells derived from a human brain show a transient transcriptional response to proinflammatory stimuli

We then performed in vitro assays to determine whether the transcriptional response observed in our TRAP data was a direct effect of LPS or IL-6 signaling on endothelial cells. An LPS challenge quickly induced IL6 expression in the immortalized microvascular brain endothelial cell line HCMEC/D3 (Figure 4A). This response was further increased if cells were challenged with LPS in the presence of the soluble form of the IL-6 receptor subunit gp80 (sIL-6Rα, denoted as LPS+R). Consistent with our previous observations in human primary endothelial cells(Martino N, et al., 2022), this co-stimulation allows for an increased transcriptional response, as evidenced by the increased SOCS3 expression. LPS+R also induced an increase in expression of multiple genes identified by our TRAP-seq (Figure 4A). This suggested an IL-6-dependent autocrine signal in response to LPS. To directly assess the effects of IL-6, we challenged these cells with a combination of IL-6 and its soluble receptor (IL-6+R). As expected, this challenge quickly induced an increase in IL6 and SOCS3 (Figure 4B). In support for a direct response to IL-6, the same challenge promoted a significant increase in the expression of CXCL10, NAMPT and CD47 (Figure 4B). We observed a similar response to LPS and IL-6 in cells originally derived from human brain cortex endothelium (HBEC-5i, Figure 5A-C). However, HCMED/D3 cells retain much stronger expression of endothelial markers than HBEC-5i cells (Figure 5D), suggesting that HBEC-5i cells lost many of its endothelial characteristics during cell culture. Notably, HCMEC/D3 demonstrated a mild but significant response to IL-6 in the absence of recombinant sIL-6Rα (Figure 4B). This is likely due to the increased endogenous sIL-6Rα expression in HCMEC/D3 cells compared to other endothelial cells (Figure 5E).

**Figure 4.**
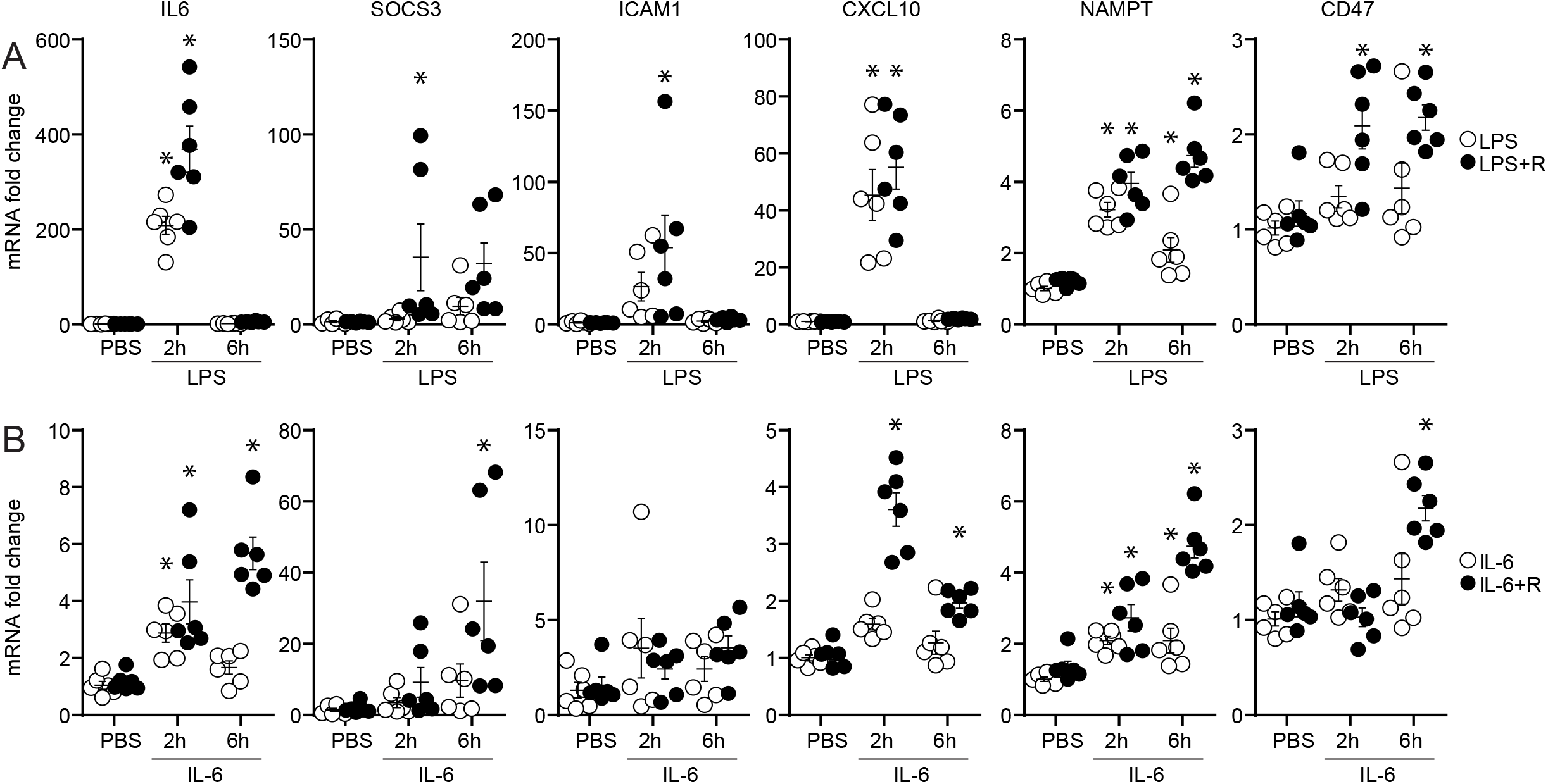
Direct transcriptional regulation by LPS and IL-6 in cultured HCMEC/D3 cells. (A) Confluent HCMEC/D3 cells were treated for two or six hours PBS, PBS+ sIL-6Rα, LPS, or LPS+sIL-6Rα (LPS+R). RT-qPCR was performed to detect gene expression levels. (B) Confluent HCMEC/D3 cells were treated for two or six hours PBS, PBS+ sIL-6Rα, IL-6, or IL-6+sIL-6Rα (IL-6+R). RT-qPCR was performed to detect gene expression levels. Asterisks, p < 0.05 (Two-way ANOVA with Sidak post-hoc test against PBS control). Data compiled from three independent experiments performed in duplicate each.

**Figure 5.**
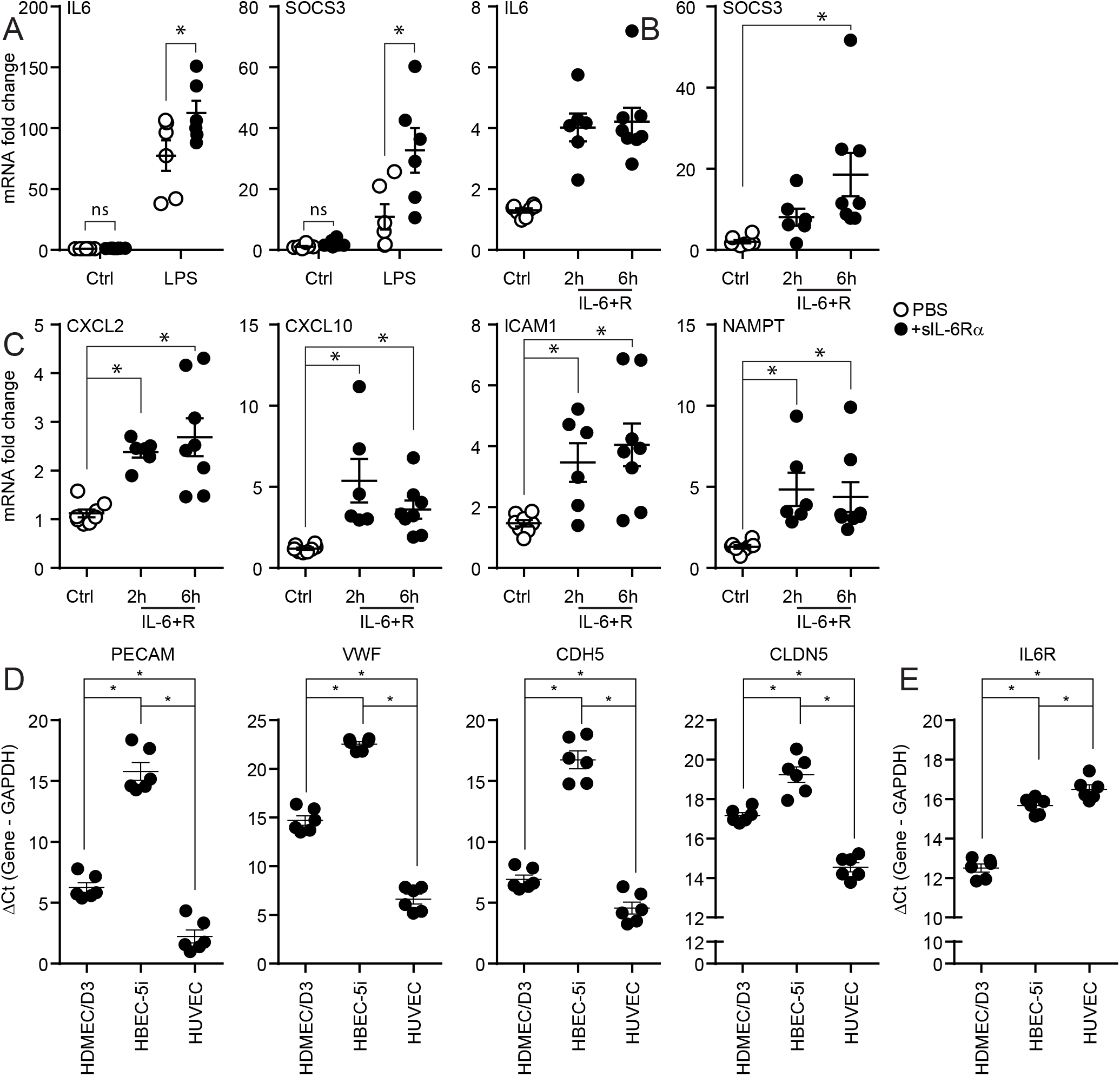
Transcriptional regulation by LPS and IL-6 in cultured HBEC-5i cells. (A) Confluent HBEC-5i cells were treated for two hours with LPS, LPS+sIL-6Rα, or PBS. RT-qPCR was performed to detect IL6 or SOCS3 mRNA levels. Two-way ANOVA with Sidak post-hoc test. (B) HBEC-5i cells were treated with IL-6+R for two or six hours (or six hours of sIL-6Rα alone as control) prior to RT-qPCR to detect IL6 or SOCS3. One-way ANOVA with Sidak post-hoc test. (C) RNA from cells treated as in (B) was used to measure CXCL2, CXCL10, ICAM1, and NAMPT mRNA levels. Asterisks, p < 0.05 (Two-way ANOVA with Sidak post-hoc test). Data compiled from three independent experiments performed in duplicate each. (D) RT-qPCR using RNA obtained from confluent HCMEC/D3 and HBEC-5i cells to probe for expression of the endothelial markers PECAM, VWF, CDH5 and CLDN5. Primary human umbilical vein endothelial cells (HUVEC) were used as positive control. Shown are ΔCt values calculated as Ct for each gene of interest minus the Ct for the housekeeping gene GAPDH. (E) RT-qPCR from cells as above to probe for expression of IL-6Rα. Asterisks, p < 0.05 (One-way ANOVA with Tukey post-hoc test). Data compiled from three independent experiments performed in duplicate each.

## Discussion

The mechanisms driving sepsis encephalopathy are thought to involve an impaired microvascular blood brain barrier, hypo- and hyper-coagulation, and reduced perfusion in response to systemic inflammatory cytokines (Crippa IA, et al., 2023;Daneman R and Prat A, 2015;Gofton TE and Young GB, 2012;Hughes CG, et al., 2016;Peng X, et al., 2021;Schramm P, et al., 2012;Sweeney MD, et al., 2019). Post-mortem analysis of septic shock decedents shows a pathology consistent with severe brain endotheliopathy, including edema, hemorrhage, microthrombi, and ischemia that is associated with higher IL-6 expression, microglia activation and neuronal apoptosis (Barichello T et al., 2022;Heming N et al., 2017;Munster BC et al., 2011;Sharshar T et al., 2004). Consistent with this mechanism, multiple experimental studies in small (Martino N, et al., 2021;Rosengarten B et al., 2007) and large (Taccone FS et al., 2014;Taccone FS et al., 2010) animals demonstrate that shock leads to severe brain microvascular dysfunction. Here, we present evidence that an acute inflammatory reaction leads to a strong but transient transcriptional profile in the brain endothelium. While this response is strongly correlated with that of the whole brain, we identified a transcriptional subset that represents an endothelial-specific response. Notably, this subset suggested an acute brain hypoxic event in response to LPS. We had previously shown that this challenge leads to a disruption in the blood-brain barrier that is worsened in mice lacking endothelial expression of the IL-6 signaling pathway inhibitor SOCS3 (Martino N, et al., 2021). We show here that loss of SOCS3 leads to a broadening of the population of genes responsive to LPS, suggesting that an overactivation of the IL-6/JAK/STAT3 pathway leads to an increased transcriptional response that could explain the severe brain injury in these mice. Notably, in WT mice, this transcriptional response returns essentially to baseline levels only days after the shock. Despite the transient nature of the response, we observed that mice that recovered from the endotoxemic shock showed mild, sex-dependent cognitive impairment, suggesting that the acute brain injury led to sustained, non-transcriptional effects.

There is strong evidence that IL-6 is a causal factor in the development of cognitive impairment (Boots EA et al., 2022;Bradburn S et al., 2017;Lyra ESNM et al., 2021;Papassotiropoulos A et al., 1999;Rothaug M et al., 2016;Wright CB et al., 2006). Much less is known about the role of IL-6 in SAE and its sequelae. A recent systematic review demonstrates increased IL-6 expression in postmortem brains of septic patients when compared to non-septic decedents (Barichello T, et al., 2022). IL-6 was found to be increased in cerebrospinal fluid (but not in plasma) of patients with all-cause encephalitis (Wang X et al., 2022). In many animal models, systemic IL-6 signaling disrupts blood-brain barrier, promotes coagulopathy, and increases leukocyte infiltration (Kang SJ and Kishimoto T, 2021). In mice, brain IL-6 is associated with aging (Godbout JP and Johnson RW, 2004) and with endotoxin-induced neuroinflammation (Henry CJ et al., 2008). Consistent with a crucial role, blockade of circulating IL-6 signaling improved survival and cognitive functions in septic (Jiang SF et al., 2023) and trauma (Hu J et al., 2018) mouse models. The transcriptional data presented here is consistent with a critical role for IL-6 signaling acting directly on the brain endothelium. LPS induced a strong expression of multiple targets of the IL-6 pathway, including SOCS3 and IL-6 itself, that was observed in both whole brain and endothelial-specific RNA isolates. Mice lacking endothelial expression of SOCS3 displayed a stronger transcriptional response to LPS. Notably, IL-6 expression was undetectable in both WT and SOCS3iEKO saline groups, but SOCS3iEKO mice showed a 4-fold stronger response than WT mice in response to LPS. Bioinformatic analysis of the transcriptional response of SOCS3iEKO mice compared to WT mice confirmed the overactivation of the JAK/STAT3 signaling axis. Moreover, direct IL-6 stimulation of cultured brain endothelial cells induced an increase in expression of multiple proinflammatory genes identified by our TRAP-seq approach. How these changes may affect brain function in shock survivors, however, remains to be determined.

Little is known regarding sex- or gender-specific risks of developing post-intensive care cognitive decline and dementia. Some clinical studies found an association with sex or gender, while others did not (Hiser SL et al., 2023;Lee M et al., 2020;Merdji H et al., 2023;Modra LJ et al., 2022;Neufeld KJ et al., 2020). Despite the endothelial and whole brain transcriptional profile returning to baseline levels after recovery from LPS, we identified mild, sex-dependent behavioral changes remaining well beyond severity scoring and temperature returned to normal levels. These findings are consistent with sex-specific cognitive deficits in mouse models of brain hypoperfusion (Gannon OJ et al., 2022;Robison LS et al., 2020;Salinero AE et al., 2020). It is not yet known however if similar processes drive cognitive decline in these two models. Future work is required to dissect the mechanisms driving this sex-specific response.

## Acknowledgements

This work was supported in part by grants R01GM124133, R01GM124133-S4 and ALZ-D-NTF 968957(APA), American Heart Association pre-doctoral award 916096 (SL), American Heart Association pre-doctoral award 908878 (AES), NINDS/NIA R01NS110749 (KLZ), and NIA U01AG072464 (KLZ).

---

**Supplemental Table 1.**

| **M&M section** | **Reagent or disposable** | **Vendor** | **Catalog Number** | **Concentration/Dose** |
| --- | --- | --- | --- | --- |
| **Mice** | Tamoxifen | Millipore Sigma | T5648 | 2 mg in 100 µL |
|  | Peanut Oil | Millipore Sigma | P2144 | N/A |
|  | Lipopolysaccharides from Escherichia coli O111:B4 (LPS) | Millipore Sigma | L4391 | 250 µg/ 250 µL |
| **Behavioral test** | Y-maze | Stoelting, Wood Dale, IL | 60180 |  |
| **Cell Culture** | Recombinant Human IL6 protein | R&D Systems | 206-IL-200/CF | 200 ng/mL |
|  | Recombinant human IL6r alpha, CF | R&D Systems | 227-SR-025/CF | 100 ng/mL |
|  | Lipopolysaccharides from Escherichia coli O111:B4 (LPS) | Millipore Sigma | L4391 | 2 µg/ml |
|  | Recombinant Human TNF-alpha | R&D Systems | 210-TA-020 | 10ng/ml |
|  | EndoGRO-MV Complete Culture Media Kit | Millipore Sigma | SCME004 | N/A |
|  | DMEM/F12 Medium | ATCC | 30-2006 | N/A |
|  | Endothelial Cell Growth Supplement | Sigma | E2759 | 40mg/mL |
|  | Fetal Bovine Serum | Sigma | 12306C |  |
|  | Penicillin-Streptomycin Solution, 100x | Corning | 30-002-CI | 1X |
|  | Gelatin | Millipore Sigma | ES-006-B | 0.1% |
|  | 0.05% Trypsin-EDTA | Gibco | 25300054 | 1x |
| **RNA isolation and RTqPCR** | PrimeScript RT Master Mix | Takara | RR036B | 1X |
|  | TRIzol reagent | Invitrogen | 15596018 | 1X |
|  | iTaq Universal SYBR Green Supermix | Bio Rad | 1725125 | 1X |
| **TRAP** | Avanti Polar Lipids, 1,2-diheptanlyl-sn-glycerol | Avanti Polar Lipids | NC9999043 | 300mM |
|  | RNAasin Plus | Promega | PAN2615 | 40 U/ml |
|  | Protein G beads | Thermo Fisher/Invitrogen | 10004D |  |
|  | Ab for TRAP | Memorial Sloan Kettering CC | HTZGFP-19C8 |  |
|  | Ab for TRAP | Memorial Sloan Kettering CC | HTZGFP-19F7 |  |
|  | HBSS no calcium, mg, phenol red | Gibco | 14175079 | 1X |
|  | Complete mini edta free easy pack | Roche | 4693159001 |  |
|  | PhosStop | Roche | 4906837001 |  |
|  | Cyclohexamide | Sigma | C1988 | 100 µg/m |
|  | DTT-biotech grade | VWR Life Sciences | 97061-340 | 0.5 mM |
|  | Potassium Chloride ACS | VWR Life Sciences | 97061-566 | 350 mM |
|  | HEPES Free Biotechnology grade | Sigma | H4034 | 1M |
|  | Magnesium Chloride | Honeywell | 63020 | 1M |
|  | Diethyl pyrocarbonate (DEPC), 5ml | Sigma | D5758 |  |
|  | Sodium Bicarbonate | Fisher Scientific | S233 | 4 mM |
|  | (D) + Glucose | Acros | 41095 |  |
|  | Igepal | MP Biomedical | CA-630 | 1X |
|  | Disposable pellet pestles with 1.5ml microtube | DWK Life Sciences | 749520-0000 |  |
|  | Cordless Pestle Motor | VWR Life Sciences | 47747-370 |  |
|  | Diamond Midi Centrifuge Tube, 5.0mL, PP, Separate Red Screw Cap, Graduated | Globe Scientific | 111580 |  |
|  | RNeasy Plus Micro Kit | Qiagen | 74034 |  |
|  | RNAse Away | Molecular Bioproducts | 7002 |  |

**Supplemental Table 2.**
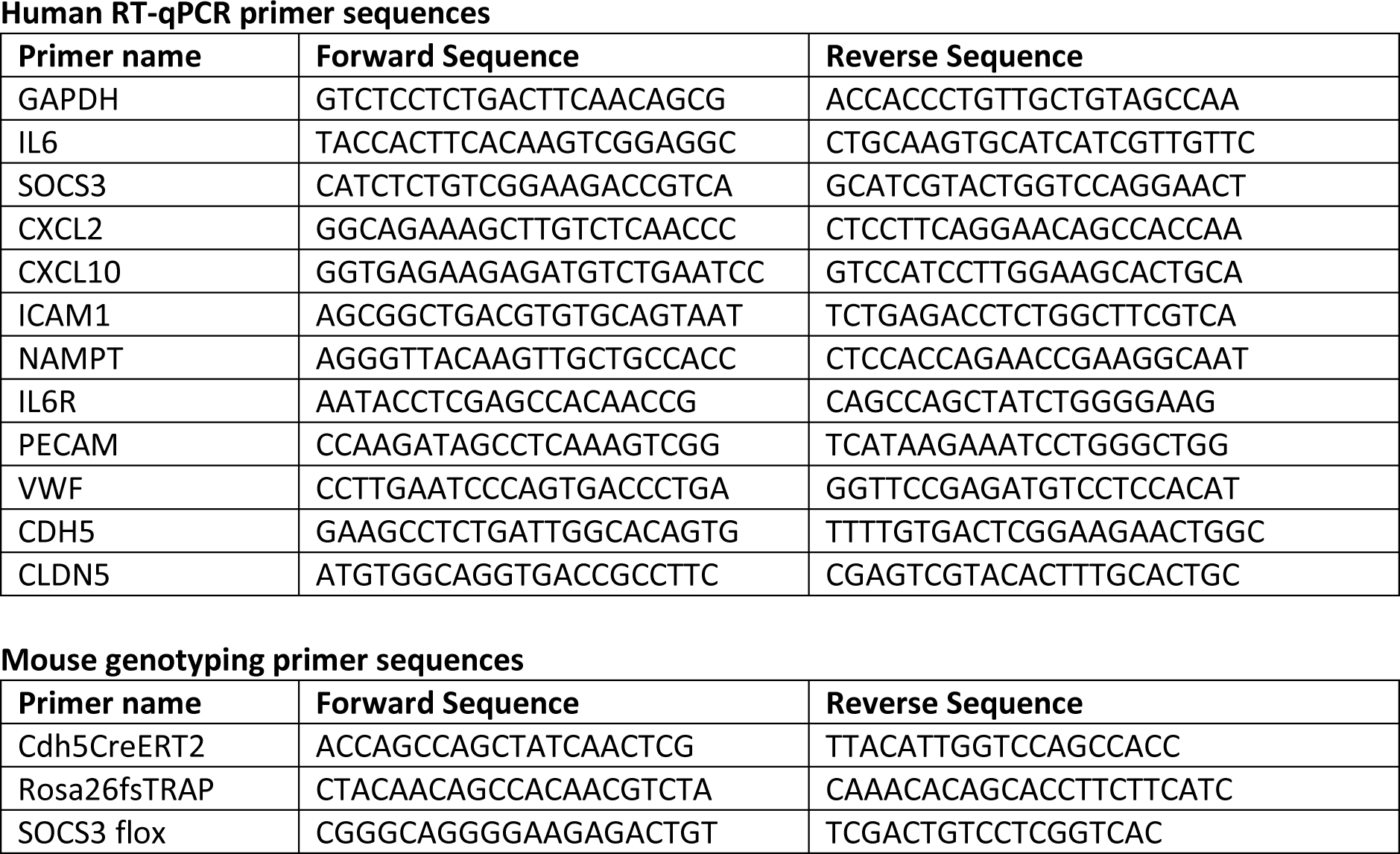

